# Low-Dose CDK4/6 Inhibitors Induce Presentation of Pathway Specific MHC ligands as Targets for Cancer Immunotherapy

**DOI:** 10.1101/2020.06.18.157800

**Authors:** Angel Charles, Christopher Bourne, Zita E. Aretz, Sung S. Mun, Tanya Korontsvit, Tao Dao, Martin G. Klatt, David A. Scheinberg

**Affiliations:** Molecular Pharmacology Program, Sloan Kettering Institute; Immunology and Microbial Pathogenesis Program, Weill Cornell Medicine; Physiology, Biophysics and Systems Biology, Weill Cornell Medicine; Pharmacology Program, Weill Cornell Medicine

## Abstract

**Purpose:** Cyclin dependent kinase 4/6 inhibitors (CDK4/6i) lead to cell-cycle arrest but also demonstrate antineoplastic activity through triggering T cell-mediated immunity. One of the potential mechanisms responsible for this immunological effect might be qualitative and quantitative changes in human leukocyte antigen (HLA) ligands on the cell surface after treatment with CDK4/6i. These changes may increase the immunogenicity of breast cancer cells offering potential synergies for combinations with cancer immunotherapies.

**Experimental Design:** We investigated the ability of two CDK4/6 inhibitors (CDK4/6i), Abemaciclib and Palbociclib, to alter the immunopeptidome at subclinical, non-toxic, levels in different breast cancer cell lines. Biochemical isolation of HLA ligands, identification by mass spectrometry and subsequent network analysis after drug treatment were used to characterize the changes in the immunopeptidome. The mechanisms for altered CDK4/6 presentation were explored.

**Results:** Low-dose treatment with 100nM of Abemaciclib and Palbociclib led to upregulation of cell surface HLA levels and induced hundreds of HLA ligands in breast cancer cell lines. These new ligands were significantly and most strongly enriched for peptides derived from proteins involved in the “G1/S phase transition of cell cycle” pathway and included among others, HLA ligands from CDK4, CDK6, Cyclin D1 and Cyclin E1. An increase in transcript, protein, and subsequent ubiquitination for Cyclin D1, which could lead to enhanced degradation of the target protein, was identified as a potential mechanism for the altered presentation of peptides.

**Conclusions:** CDK4/6i treatment gave rise to drug-induced antigens through cell cycle disruption and increased antigen presentation. Interestingly, these induced HLA ligands are often sourced from the proteins of the CDK4/CDK6/CCND1 complex or more downstream interaction partners, providing evidence that inhibition of a distinct cellular pathway leads to increased presentation of the proteins involved. These findings suggested CDK4/6i provided a tool for highly selective induction of HLA ligands that may be targeted by T cell-based immunotherapeutics.

**Translational Relevance:** These data demonstrated that low-dose treatment of breast cancer cells with CDK4/6 inhibitors, Abemaciclib and Palbociclib, induced marked changes in presentation of HLA ligands, especially from proteins involved in the G1/S phase transition, the phase in which these drugs arrest the cells. Enhanced ubiquitination and degradation was identified as a mechanism for the altered presentation for one of the relevant proteins. The induced HLA ligands may provide ideal specific targets for combination immunotherapies. The data show for the first time that selective inhibition of a distinct pathway can lead to specific presentation of HLA ligands in breast cancer cells. This work supports the rationale for testing the combination of low-dose CDK4/6i with immunotherapeutic agents, such as immune checkpoint blockade antibodies or T-cell-based approaches specifically directed against one of the induced HLA ligands.

## Introduction

The development of targeted cancer therapies is an important part of cancer therapy now, as traditional antineoplastic agents (e.g. chemotherapy, radiation) are non-specific, and prone to off-target toxicities to healthy tissues (1,2). Successful therapies such as checkpoint blockade and adoptive T-cell therapy target cancer-associated antigens (3,4), but are still non-specific immune activators associated with severe toxicities. Checkpoint blockade therapies rely on the ability of T-cells to recognize and kill cells that express cognate HLA/peptide complexes presenting neoantigens (5,6).

Mutated neoantigens as HLA ligands provide ideal cancer specificity, but are patient-specific (7). Therefore, more broadly expressed tumor-selective targets are needed to develop more widely applicable immunotherapies. T-cell receptor (TCR)-based therapies, including T cells and checkpoint blockade inhibitor antibodies, can be directed to proteins expressed inside the cell when presented as peptides on the surface, theoretically making most proteins in the cell potential targets (8,9). This key feature of TCR surveillance can be exploited to target intracellular, oncogenic and tumor selective pathways.

Classic cancer neoepitopes are derived from mutations (6). However, drugs that influence oncogenic signaling pathways can lead to the presentation of HLA ligands on cancer cells that are usually not displayed in healthy tissues (10,11). These drug-induced HLA ligands may be immunogenic. For example, inhibition of the mitogen-activated protein kinase (MAPK) pathway in cancers in which this pathway is activated has been shown to lead to improved peptide/MHC target recognition and killing by T cells and TCR-mimic antibodies (10). Additionally, the inhibition of two tyrosine kinases, anaplastic lymphoma kinase (ALK), and rearranged during transfection (RET), in the cancers in which these pathways are oncogenic, promotes HLA class I antigen presentation, and induces expression of new peptides that are recognized by T cells (11).

Despite rapid advancements in the treatment of breast cancer, many patients still relapse and are incurable. Immunotherapy for breast cancer has been largely unsuccessful (12). Therefore, discovery of therapeutic strategies that would increase susceptibility of breast cancer to immunotherapies are highly attractive. Cyclin dependent kinase 4/6 inhibitors (CDK4/6i), which prevent the transition from G1 to the S phase of the cell cycle, have been shown to increase the expression of HLA class I molecules in multiple breast cancer cells lines (13,14). CDK4/6i treatment has also been demonstrated to improve T cell infiltration into the tumor, and to work synergistically with immune checkpoint blockade (e.g. PD-L1)(13).

We hypothesized that the increase in antigen presentation by CDK4/6 inhibitors would lead to presentation of new, drug-induced cancer-associated antigens that might be useful targets of immunotherapies. Utilizing HLA-peptide immunoprecipitation in conjunction with mass spectrometry (MS), we identified the presentation of HLA ligands derived from proteins involved in G1/S phase cell cycle transition in which cells were treated with CDK4/6i. Many of these drug-induced HLA ligands had not been identified in healthy tissues, which renders them appealing targets for TCR-based therapies and TCRmimic antibodies (9). Moreover, this concept demonstrated a small molecule drug’s disruption of a molecular pathway led to increased presentation of that particular complex and pathway and illustrates how CDK4/6 inhibitors could potentially synergize therapeutically with immunotherapeutic agents.

## Materials and Methods

### Human Cell Lines

MCF7 (HLA-A*02:01, HLA-B*18:01, HLA-B*44:02, HLA-C*05:01) and T47D (HLA-A*33:01, HLA-B*14:02, HLA-C*08:02) cell lines were maintained were cultured in RPMI medium (Gibco) supplemented with 10% FBS (Gibco), penicillin and streptomycin, and 1% L-glutamate. All cell lines were obtained from ATCC, and tested negative for mycoplasma.

### Antibodies and commercial reagents

Antibodies used in this study follow: APC anti-Human HLA-A, B, C (clone W6/32) from Biolegend, FITC Anti-Human HLA-A2 from BD Biosciences, were used for flow cytometry. Unlabeled clone W6/32 antibody from BioXcell, was used for HLA class I/peptide immunoprecipitation for mass spectrometry. Anti-Human Cyclin D1 (ab134175) from Abcam, and Anti-Multi ubiquitin magnetic beads from MBL International, were used for Western blot. All antibodies were kept at +4C. Abemaciclib and Palbociclib were purchased from Sellekchem. The drugs were diluted to 10mM stocks in DMSO, and kept at −20C.

### Dose Response relationships with drugs

15,000 cells were cultured in 96 well plates with serial dilutions of Abemaciclib and Palbociclib. After 4 days, cells were resuspended in Cell Titer-Glo Luminescent Cell Viability Assay from Promega for 10 minutes, and agitated at room temperature. Luminescence in the 96 well plates was read using a SpectroMax2 (Molecular Devices, Sunnyvale, CA, USA).

### Flow Cytometry

Cells were treated with 100nM Abemaciclib or Palbociclib for 1, 3, 5 or 7 days. Cells were harvested, washed with PBS, and labeled with 1:50 dilution of APC Anti-Human HLA-ABC and FITC Anti-Human HLA-A2 in buffer (2% FBS, 0.1% sodium azide, in PBS) for 30 minutes. Cells were analyzed using a Fortessa Flow Cytometer from BD Biosciences or a Guava flow cytometer from Millipore.

### Quantitative reverse-transcriptase PCR

Cells were treated with 100nM Abemaciclib or Palbociclib for 1, 3, 5 or 7 days. Cells were harvested, washed with PBS, and RNA was extracted using Qiagen RNA Easy Plus (Qiagen; #74134). cDNA was created using qScript cDNA SuperMix (Quantabio; #95048) according to recommended cycling times and temperatures in thermocycler. qPCR was performed using PerfeCTa FastMix II (Quantabio; #95118) and TaqMan real-time probes purchased from Life Technologies: HLA-A (Hs01058806_g1), beta-2-microglobulin (Hs00187842_m1), TAP1 (Hs00388677_m1), and TAP2 (Hs00241060_m1), Cyclin D1 (Hs00765553_m1), GAPDH (Hs02758991), Actin B (Hs99999903_m1). Data were normalized to baseline expression of GAPDH or Actin B(11).

### Immunopurification of HLA class I ligands

Immunopurification affinity columns were prepared in the lab. 40 mg of Cyanogen bromide-activated-Sepharose 4B (Sigma-Aldrich, cat. # C9142) was activated with 1 mmol/L hydrochloric acid (Sigma-Aldrich, cat. # 320331) for 30 minutes. Subsequently, 0.5 mg of W6/32 antibody (Bio X Cell, BE0079; RRID: AB_1107730) was coupled to sepharose in the presence of binding buffer (150 mmol/L sodium chloride, 50 mmol/L sodium bicarbonate, pH 8.3; sodium chloride: Sigma-Aldrich, cat. # S9888, sodium bicarbonate: SigmaAldrich, cat. #S6014) for at least 2 hours at room temperature. Sepharose was blocked for 1 hour with glycine (Sigma-Aldrich, cat. # 410225). Columns were equilibrated with PBS for 10 minutes. MCF7 and T47D breast cancer cells were treated with DMSO, 100 nmol/L Abemaciclib, or 100 nmol/L Palbociclib for seven days. Cells (1×107 *per condition*) were harvested with Cellstripper (Corning, cat. # 25-056-CI) and washed three times in ice-cold sterile PBS (Media preparation facility MSKCC). Afterward, cells were lysed in 1 mL 1% CHAPS (Sigma-Aldrich, cat. # C3023) in PBS, supplemented with 1 tablet of protease inhibitors (Complete, cat. # 11836145001) for 1 hour at 4C. This lysate was spun down for 1 hour at 20,000 g at 4C. Supernatant was run over the affinity column through peristaltic pumps at 1 mL/minute overnight at 4C. Affinity columns were washed with PBS for 15 minutes, run dry, and HLA complexes subsequently eluted three times with 200 mL 1% trifluoracetic acid (TFA, Sigma/Aldrich, cat. # 02031). For the separation of HLA ligands from their HLA complexes, tC18 columns (Sep-Pak tC18 1 cc VacCartridge, 100 mg Sorbent per Cartridge, 37–55 mm Particle Size, Waters, cat. # WAT036820) were prewashed with 80% acetonitrile (ACN, Sigma-Aldrich, cat. # 34998) in 0.1% TFA and equilibrated with two washes of 0.1% TFA. Samples were loaded, washed again with 0.1% TFA, and eluted in 400 mL 30% ACN in 0.1%TFA followed by 400 mL 40% ACN in 0.1%TFA, then 400 mL 50% ACN in 0.1%TFA. Sample volume was reduced by vacuum centrifugation for mass spectrometry analysis (11).

### LC-MS/MS analysis of HLA ligands

Samples were analyzed by a high-resolution/high-accuracy LCMS/MS (Lumos Fusion, Thermo Fisher). Peptides were desalted using ZipTips (Sigma Millipore; cat. #ZTC18S008) according to the manufacturer’s instructions and concentrated using vacuum centrifugation prior to being separated using direct loading onto a packedin-emitter C18 column (75 mm ID/12 cm, 3 mm particles, Nikkyo Technos Co., Ltd). The gradient was delivered at 300 nL/ minute increasing linear from 2% Buffer B (0.1% formic acid in 80% acetonitrile)/98% Buffer A (0.1% formic acid) to 30% Buffer B/70% Buffer A, over 70 minutes. MS and MS/MS were operated at resolutions of 60,000 and 30,000, respectively. Only charge states 1, 2, and 3 were allowed. 1.6 Th was chosen as the isolation window and the collision energy was set at 30%. For MS/MS, the maximum injection time was 100 ms with an AGC of 50,000 (11).

### Mass spectrometry data processing

Mass spectrometry data were processed using Byonic software (version 2.7.84, Protein Metrics) through a custom-built computer server equipped with 4 Intel Xeon E5-4620 8-core CPUs operating at 2.2 GHz, and 512 GB physical memory (Exxact Corporation). Mass accuracy for MS1 was set to 6 ppm and to 20 ppm for MS2, respectively. Digestion specificity was defined as unspecific and only precursors with charges 1, 2, and 3, and up to 2 kDa were allowed. Protein FDR was disabled to allow complete assessment of potential peptide identifications. Oxidization of methionine, N-terminal acetylation, phosphorylation of serine, threonine, and tyrosine were set as variable modifications for all samples. All samples were searched against the UniProt Human Reviewed Database (20,349 entries, http://www.uniprot.org, downloaded June 2017). Peptides were selected with a minimal log prob value of 2 resulting in a 1% false discovery rate and were HLA assigned by netMHC 4.0 with a 2% rank cutoff (11).

## Results

### Immunologic activity of CDK4/6i on breast cancer cell lines

Abemaciclib and Palbociclib are a class of small molecule inhibitors of both CDK4 and CDK6 that are FDA approved for the treatment of hormone receptor positive (HR+) breast cancer in post-menopausal women (15). We treated MCF7 and T47D breast cancer cell lines with abemaciclib and palbociclib, respectively. First, we aimed to determine the lowest dose of CDK4/6i that would upregulate cell surface HLA expression. MCF7 and T47D were treated with varying concentrations of abemaciclib and palbociclib, respectively for 7 days. Flow cytometric analysis showed 100 nM as the lowest dose where HLA was significantly upregulated in both cell lines (data not shown). Importantly, 100nM of either drug was about 100 fold lower than the respective IC50 values for killing cells (Suppl Fig 1B-C), and 2 to 3 fold lower than plasma concentrations in treated patients (16). Therefore, observed effects were unlikely to be attributed to cell death processes. By bright field microscopy and flow cytometry, the 100nM dose of Abemaciclib resulted in an increase in MCF7 cell size when compared to DMSO treatment (Fig1A, Suppl Fig 1C), at the same time point, T47D cells also show an increase in overall cell size, take on a more spherical morphology, and increase in granularity when compared to DMSO treated cells (Fig 1C, Suppl Fig 1D).

**Figure 1.**
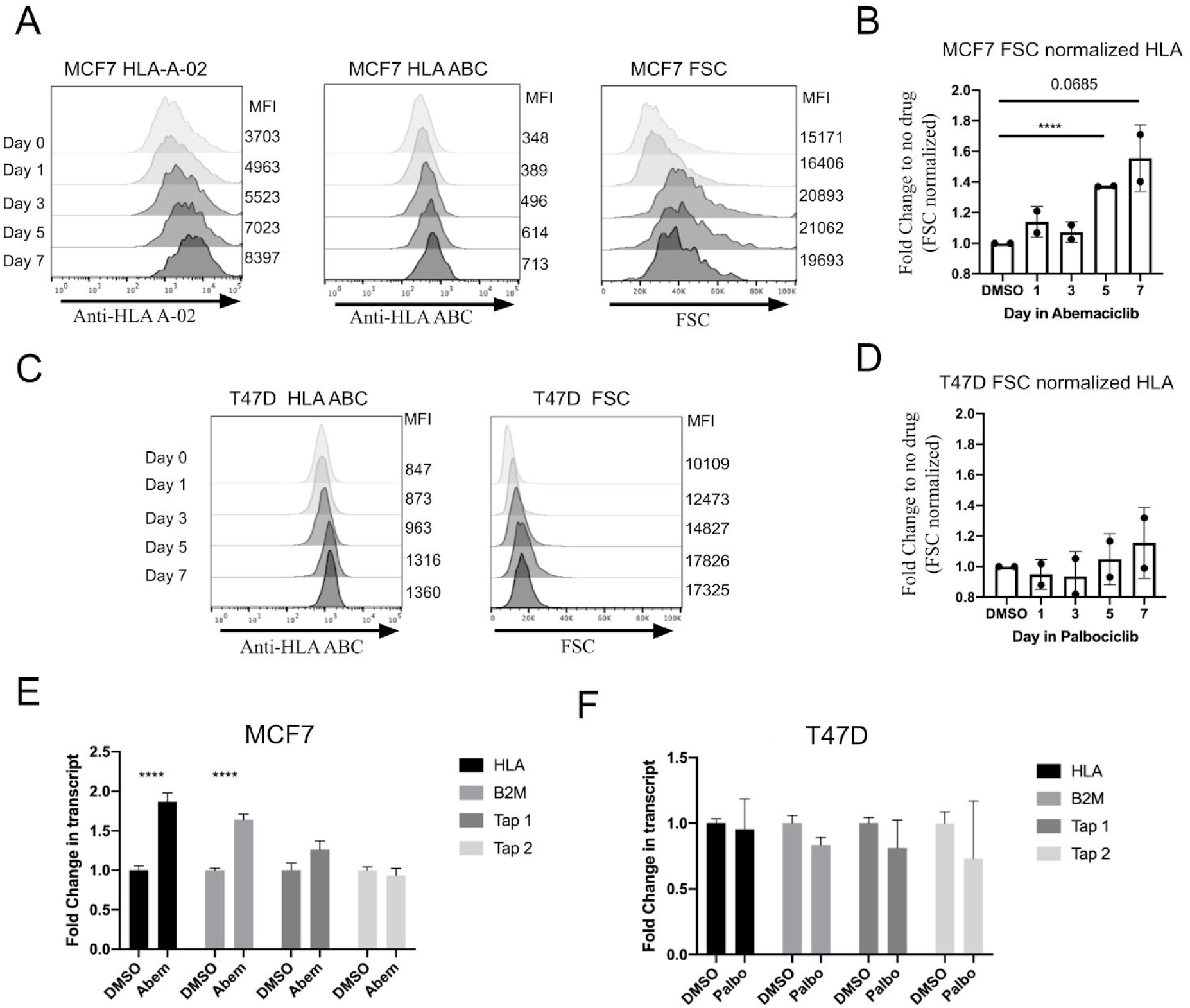
CDK4/6 inhibitors increase HLA surface expression. (A) MCF7 breast cancer cells were treated with 100 nM Abemaciclib for indicated times in the Y axis. Flow cytometry for HLA-A02 levels (left), pan HLA levels (middle), and forward scatter (right) are displayed against corresponding mean fluorescence intensity (MFI) in X axis. The quantification is annotated outside on the right side. (B) Fold change in MFI from panel A. HLA expression was normalized to forward scatter (FSC) to account for cell size changes and expressed as fold change to DMSO. (C) T47D breast cancer cells were treated with 100nM Palbociclib for indicated times using flow cytometry as in panel A. (D) Fold change in MFI from panel C. HLA expression was normalized to forward scatter (FSC) to account for cell size changes and expressed as fold change to DMSO. (E-F) qRT-PCR for key protein involved in antigen presentation transcripts noted on insert legend was performed on (E) MCF7 cells and (F) T47D cells, 7 days after treatment with a single dose of 100nM abemaciclib or palbociclib on day 1.

### CDK4/6i increases HLA class I surface expression and cell size in breast cancer cells

CDK4/6i are known to upregulate HLA class I ligand expression in breast cancer cell lines. In a 7-day time course, low-dose (100nM) Abemaciclib treatment resulted in an increase in HLA-A02 and pan-HLA class I surface expression per cell (Figure 1A, left and middle), as well as overall cell size in MCF7 breast cancer cells (Figure 1A, right). When pan-HLA class I surface expression was recalculated to account for the increase in cell size, the treatment still showed a 50% increase in pan-HLA class I surface expression density in MCF7 cells treated with Abemaciclib compared to DMSO by day 7 (Figure 1B). These results suggest that HLA class I surface expression is increased independently of the increase in cell surface size in MCF7 cells. Sub-therapeutic dosing of Palbociclib also caused an increase in pan-HLA class I surface expression per cell, as well as cell size in T47D breast cancer cells (Figure 1C). However, when the HLA class I expression was normalized to cell size, there was no increase in HLA class I surface density (Figure 1D). These results suggested that Palbociclib increased T47D HLA class I surface expression largely through increases in cell size.

The increase in HLA class I surface expression that was seen with CDK4/6i treatment could be due to a number of factors including increased transcription or translation of proteins involved in HLA expression, or increased stabilization of HLA molecules on the cell surface (11). In order to gain insight into the mechanism underlying the observed phenomena, we performed qRT-PCR analysis on key gene products involved in antigen presentation. In MCF7 cells treated with 100nM of Abemaciclib, there was a 1.5-2 fold increase in the transcript levels of both HLA and β-2 Microglobulin (b2M), but little change in transcript levels of Transporter Associated with Antigen Processing 1 and 2 (TAP1 and TAP2) when compared to DMSO (Figure 1E). Moreover, in T47D cells treated with Palbociclib, there were no significant changes in HLA, b2M, or TAP1 transcript levels, and a slight decrease in TAP2 transcript levels when compared to DMSO (Figure 1F). These results imply that the increase in HLA class I surface protein levels observed in CDK4/6i treated breast cancer cells were not consistent with a broad upregulation in antigen presentation, which differs from previous observations with MAPK inhibitors (10,11).

### Mass spectrometry identified changes in the immunopeptidome after CDK4/6i treatment

As the immunopeptidome of breast cancer cells after CDK4/6i treatment has not been studied yet and changes in the HLA levels on the surface suggested a potential alteration in the presented repertoire, we biochemically isolated HLA class I peptides from breast cancer cells treated with either Abemaciclib, Palbociclib, or DMSO, assigned them to their HLA alleles through prediction algorithms and then mapped the peptides to their proteins of origin. The HLA class I alleles expressed in the MCF7 cell line are HLA-A*02:01, B*18:01, B*44:02, C*05:01; in the T47D cell line the alleles are A*33:01, B*14:02, C*08:02 (17).

In the MCF7 cell line, CDK4/6i treatment increased the number of uniquely identified HLA ligands 1.5 or 1.9-fold for Palbociclib or Abemaciclib, respectively resulting in 160 or 200 unique HLA class I ligands (Fig. 2A, B). Compared to DMSO both drugs taken together induced the presentation of more than 200 new HLA class I ligands at concentrations of 100nM of either drug (Fig. 2C). Further analysis of the HLA ligands showed the increase in HLA ligand presentation changed neither the allelic nor the length distribution of the HLA ligands demonstrating consistency of the approach and excluding potential bias through different processing after drug treatment (Suppl. Fig. 2A, B). In the MCF7 cell line, HLA-A*02:01, and HLA-C*05:01 loci presented more peptides in the CDK4/6i treatment groups compared to DMSO. Fewer peptides from the HLA-B*18:01 and HLA-B*44:02 were found in any treatment group, with no significant change in the peptide quantity or length in the CDK4/6i groups compared to DMSO (Suppl. Fig. 2A, B).

**Figure 2.**
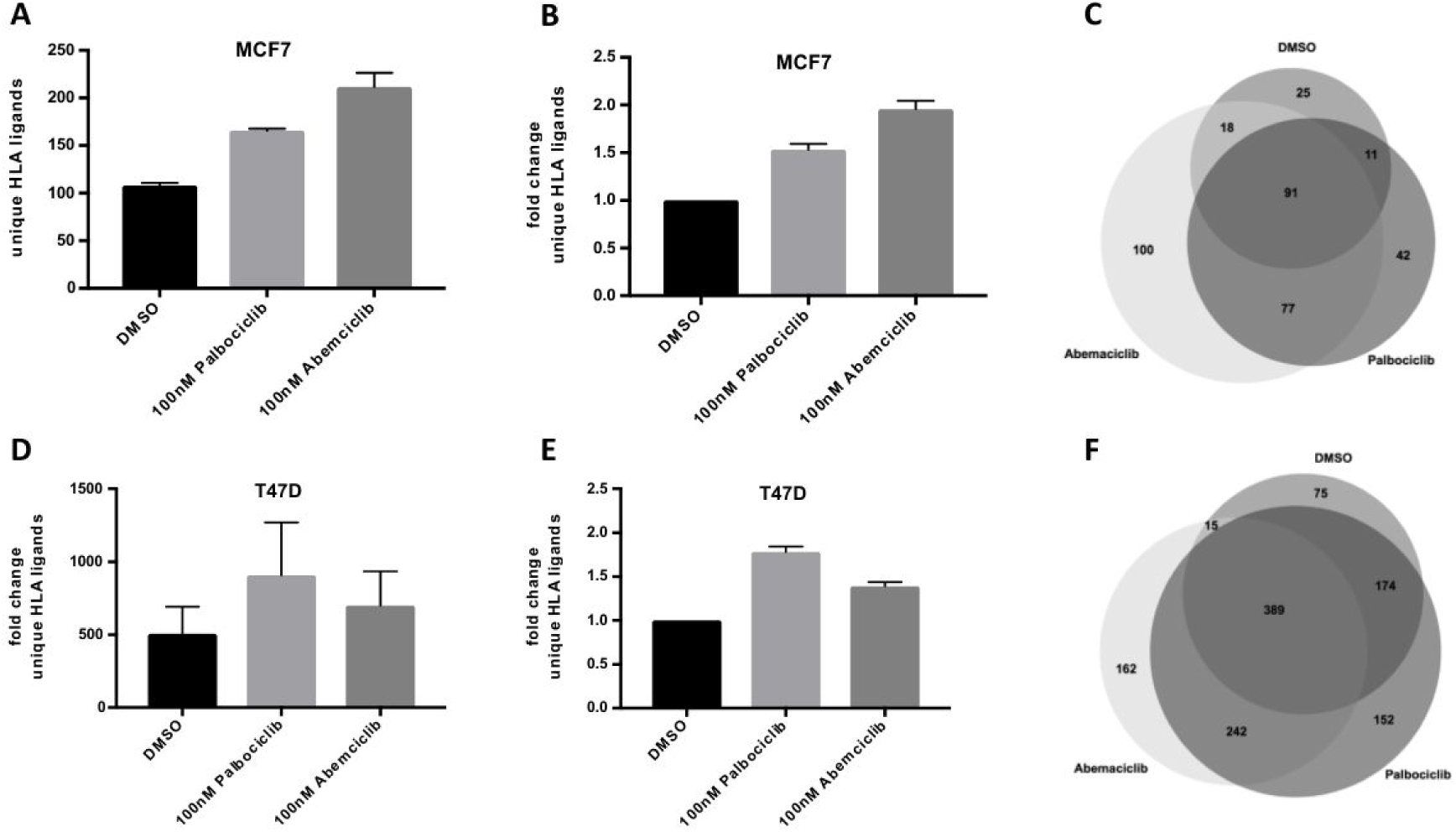
CDK4/6i induce changes in the immunopeptidome. Immunoprecipitation of HLA complexes and subsequent isolation and analysis of HLA-bound peptides by LC-MS/MS demonstrate qualitative and quantitative changes in the immunopeptidome after 7-day treatment with either 100nM DMSO, Abemaciclib or Palbociclib. MCF and T47D cells, respectively, showed absolute increase in unique HLA ligands (A, D), and relative changes of unique HLA ligands normalized to DMSO (B, E). Venn diagrams of the mean of total unique ligands from two biological replicates per treatment condition were displayed (C, F Error bars indicate SD.)

In the T47D cell line much higher total numbers of HLA ligands were detected consistent with previously published data that determined that absolute HLA transcript levels are almost 3-fold higher in T47D compared to MCF7 cells (18). With an average of 908 and 700 unique HLA ligands identified in the Palbociclib and Abemaciclib treated cell lines compared to 506 in DMSO treated cells the relative increase was about 1.7 and 1.4-fold respectively (Fig. 2 D, E). This resulted in a total of more than 500 new HLA class I ligands induced by drug CDK4/6i treatment (Fig. 2F). Again, no changes in the distribution of allelic preference or length were detected but HLA ligands were more equally distributed between the different HLA alleles (Suppl. Fig. 1C, D).

### CDK4/6i-induced HLA ligands are derived from proteins enriched in G1/S cell cycle transition

To further understand if there is a correlation between the inhibition of the CDK4/6 complex and the change in the immunopeptidome, we analyzed the source proteins of the drug-induced HLA ligands. Results from biological replicates of the different treatment conditions were combined for every cell line individually and source protein lists were generated from either all proteins in the DMSO samples or from all source proteins induced from CDK4/6i treatment (Fig. 3 A, B). The network analysis was focused on physical interactions rather than co-expression or genetic interactions, to address the direct physical inhibition of the CDK4/6 complex and pathways by this drug class. Network analyses clearly demonstrated an enrichment for proteins from the G1/S transition of cell cycle. The enrichment of this class of proteins was not only the top hit for both cell lines after drug treatment, but also was not enriched in the cells after DMSO-treatment, suggesting a drug-specific effect that is consistent with the known mechanism of CDK4/6i leading to cell cycle arrest in G1 phase (Fig. 3 A, B). Building a network analysis of all the proteins identified in our study through CDK4/6i induction that are involved in the “G1/S transition of mitotic cell cycle” illustrated three subgroups relevant to the process: the CDK4/6/CCND1-complex (Fig. 3C, lower left)), the mcm-complex (Fig. 3C, middle) and the proteasome (Fig. 3C, upper right). Overall, a total of 60 unique HLA ligands (where post-translationally modified peptides are considered distinct from the unmodified counterpart) from various alleles and both cell lines were exclusively detected after drug treatment. These peptides were all derived from proteins from the inhibited pathway (Table 1). Of note, HLA ligands were repeatedly identified from the directly inhibited molecules CDK4 and CDK6, from the main interacting partner CCND1, and also from more distant physical interactors in the G1/S transition pathway, e.g. components of the mcm-complex. Other important proteins specifically induced after CDK4/6i treatment included the prognosis relevant proteins cyclin E1 (CCNE1) and the G1 to S phase transition protein 1 homolog (GSPT1).

**Table 1.**
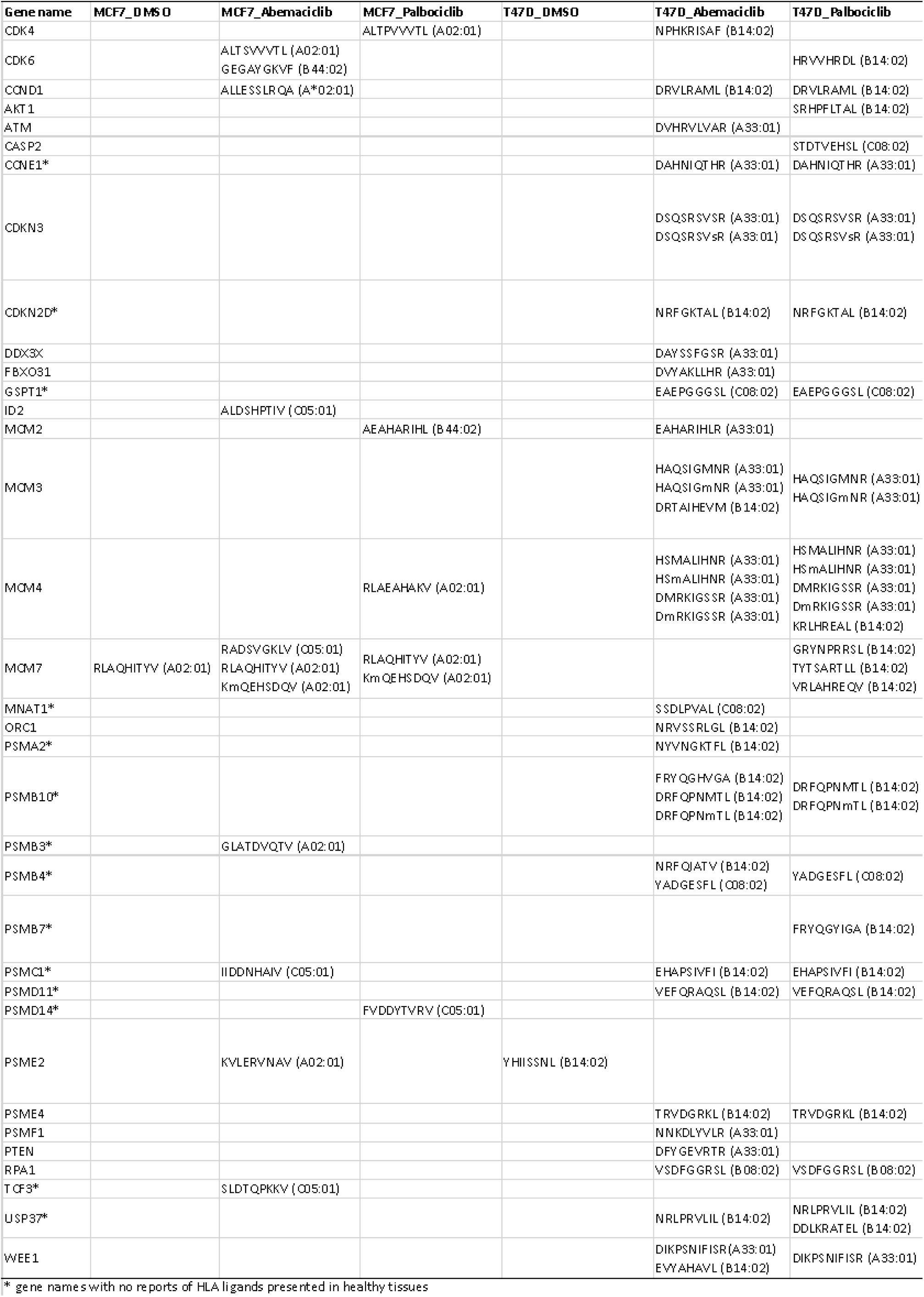
CDK4/6i induced peptides derived from prote ins contributing to the Go term G1/s transtion of mitotic cell cycle

**Figure 3.**
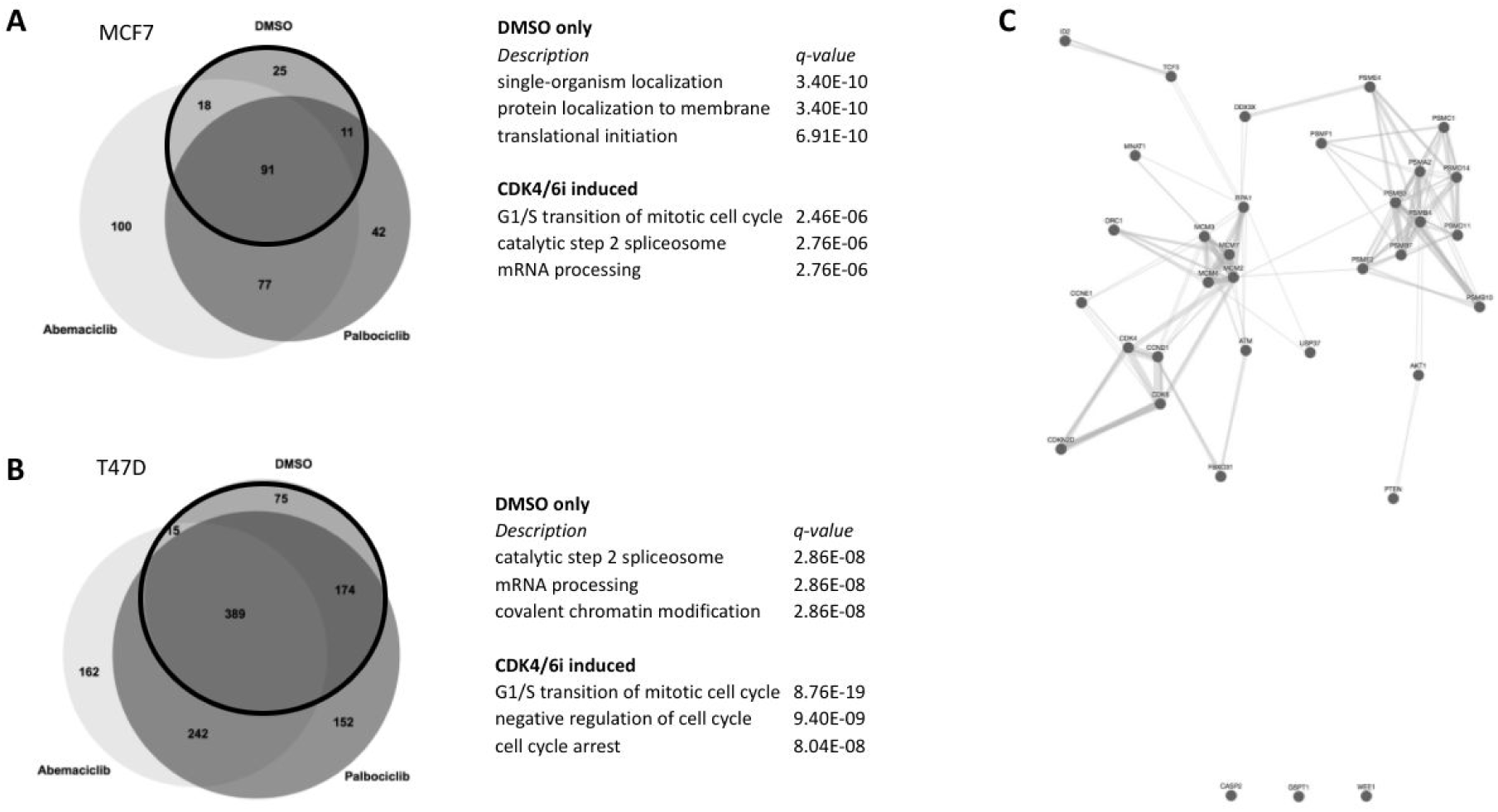
CDK4/6i induce MHC ligands enriched from source proteins contributing to “G1/S transition of mitotic cell cycle”. Source proteins from peptide sequences detected either in the DMSO sample (inside the bold lined circle) or source proteins from CDK4/6i induced peptides (outside the bold lines circle) were used for network analyses. Top 3 enriched GO terms are shown for MCF7 cells (A) and T47D cells (B). (C) Network analysis of physical interactions from all proteins induced by CDK4/6i treatment contributing to the GO term “G1/S transition of mitotic cell cycle”.

### Increased degradation of pathway specific source proteins as potential mechanism for increased presentation as HLA ligands

In order to better understand the mechanism underlying the induction of so many HLA ligands derived from the inhibited pathway we focused on CCND1 as an important target as it is important for carcinogenesis in many cancer types (15,16) and proven to lead to immunogenic HLA ligands. Furthermore, CCND1 is the direct interacting partner of CDK4/6 (16) and we exclusively identified in both cell lines, peptides derived from CCND1 after CDK4/6i treatment (ALLESSLRQA in complex with HLA-A*02:01 on MCF7 cells and DRVLRAML in complex with B*14:02 on T47D cells). We first evaluated the transcript and protein expression levels of cyclin D1 in CDK4/6i treated cells compared to DMSO during a 7-day time course. As anticipated, we observed compensatory upregulation of Cyclin D1 transcript levels in both MCF7 and T47D cells treated with CDK4/6i at days 5 and 7 (Figure 4A). Western blot analysis of the protein expression levels of Cyclin D1 in both breast cancer cell lines largely paralleled that of transcript, peaking at days 5 and 7 (Figure 4C). This indicated that at the protein level, increased Cyclin D1 expression correlated with the increase in Cyclin D1 peptide presentation in CDK4/6i treated breast cancer cells, suggesting that inhibition of this pathway might lead to a compensatory increase in protein levels and antigen presentation.

**Figure 4.**
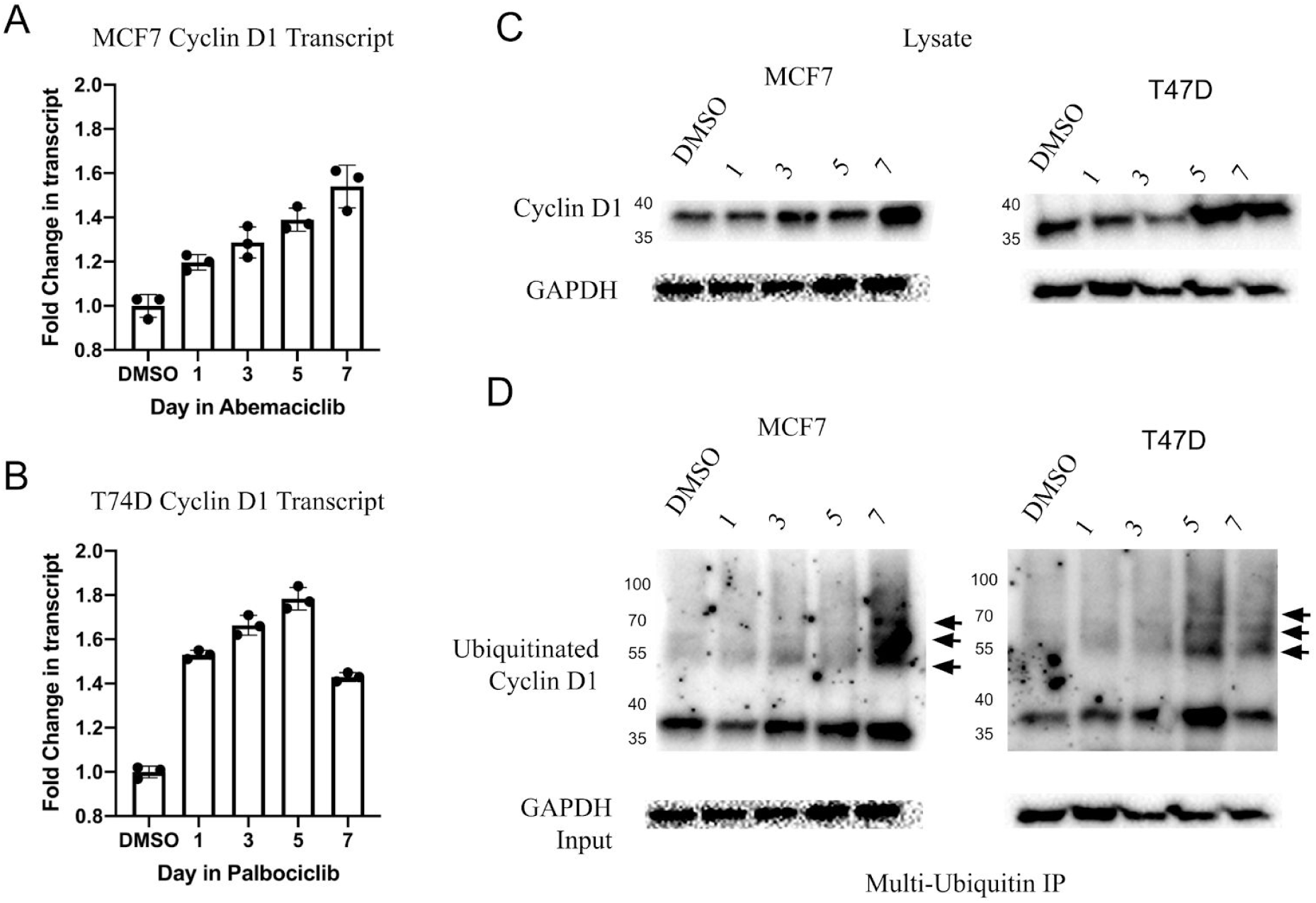
CDK4/6i alter cell cycle pathway for immunotherapeutic targeting. (A-B) MCF7 and T47D breast cancer cells were treated with 100 nM abemaciclib or palbociclib, respectively, for indicated time series. qRT-PCR of Cyclin D1 compared to Actin B was normalized to DMSO treated cells. (C) MCF7 and T47D cells were treated as described in (A). Cell lysates were Western blotted for cyclin D1. Molecular weight is provided on the left axis(kD). (D) Ubiquitinated proteins were immunoprecipitated from cell lysates in (C), and Western blot was performed for ubiquitinated Cyclin D1. Arrowheads denote higher molecular weight bands of Cyclin D1, indicative of increased ubiquitination. Molecular weight is provided on the left axis (kd).

Next, we sought to understand the mechanism whereby increased Cyclin D1 was being presented more in cells treated with CDK4/6i. Proteins degraded through the multi-ubiquitin pathway can be presented on HLA (19). We immunoprecipitated multi-ubiquitinated proteins and then performed a western blot for Cyclin D1 over time. The amount of ubiquitinated cyclin D1 also increased in CDK4/6i treated MCF7 and T47D cells compared to DMSO (Figure 4D). The increased ubiquitination of Cyclin D1 was further evidenced by higher molecular weight bands at later time points. Taken together, these results support a model whereby CDK4/6i led to compensatory upregulation of cell cycle proteins, followed by their degradation and presentation on HLA (Figure 5).

**Figure 5.**
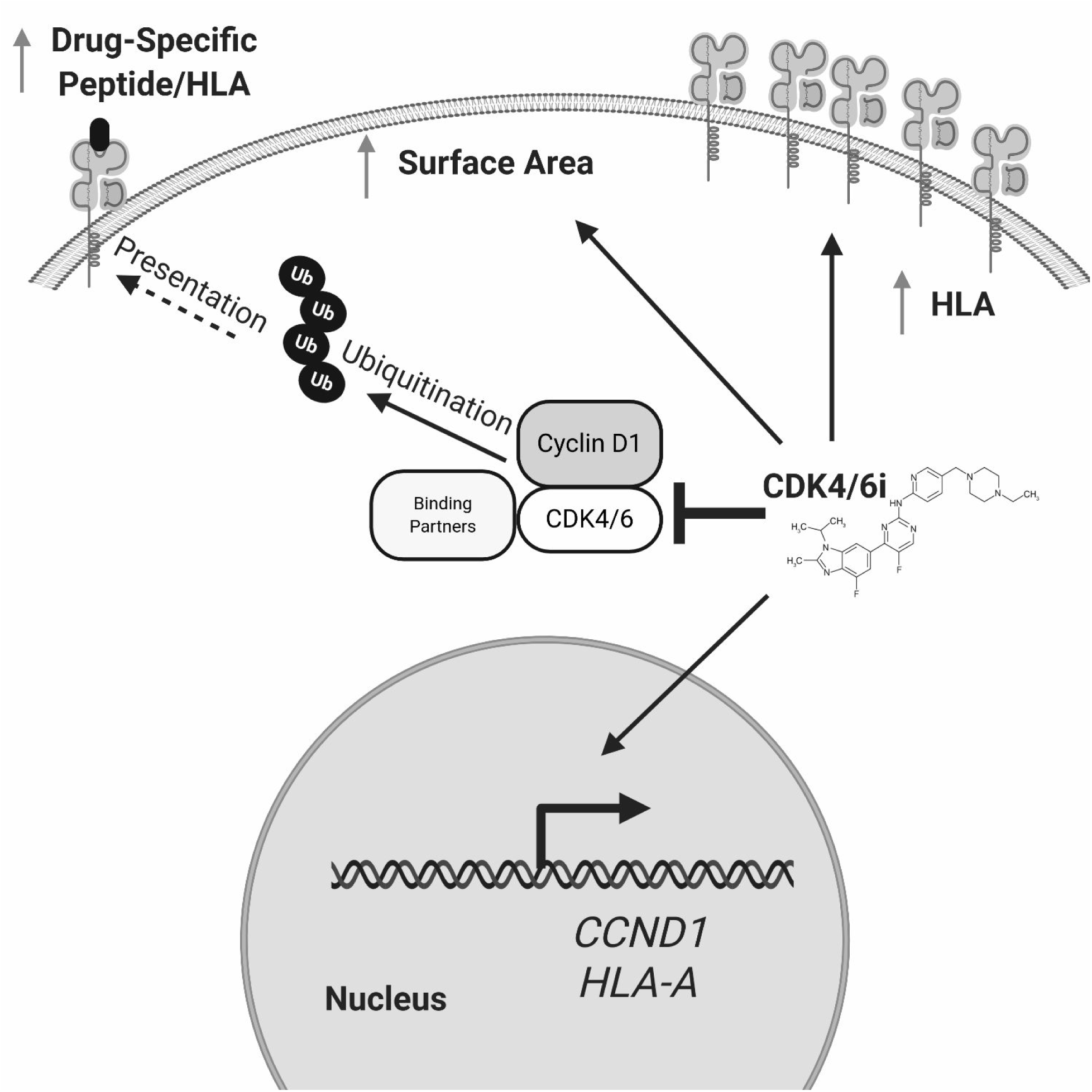
Diagram of proposed mechanism. CDK4/6i inhibit CDK4/6 and their binding partners, leading to ubiquitination. CDK4/6i also promotes compensatory upregulation of cell cycle proteins, HLA, and increased surface area. Together, these mechanisms lead to presentation of a wide variety of cancer-specific peptides derived from cell cycle effectors.

## Discussion

Immunotherapies, such as immune checkpoint blockade therapy are now standard of care for many cancer types (20). However, for many cancer types, checkpoint blockade remains unsuccessful likely due to lack of presentation of suitable immunogenic epitopes, e.g. neoepitopes or tumor associated antigens on the cancer cell surface (21,22). Similarly, adoptive T cell therapies, ImmTacs, vaccines, or TCRmimic antibodies also depend on the reliable presentation of the HLA ligand on the target cells which they are specific for (7,9,23,24).

Here we demonstrate that non-toxic doses (100nM) of CDK4/6 inhibitors Abemaciclib and Palbociclib can induce large drug-induced changes in HLA class I antigen presentation in breast cancer cells, potentially making them more susceptible to immune mediated killing by CD8+ T cells. CDK4/6 inhibitors led to an increase in total surface area of treated breast cancer cells, alongside a transcriptional upregulation of HLA in the MCF7 cell line. Although increases in HLA may for the most part be due to enlarged cell volume in the T47D cell line, we still observed quantitative and qualitative changes in the immunopeptidome by mass spectrometry in both the MCF7 and T47D cell lines. These results indicated that an increase in HLA complex levels may broaden the repertoire of detectable presented antigens even if higher HLA cell surface levels are due to an expanded surface area. In contrast to previous results, we did not observe significant changes in transcription of antigen presentation machinery, as a whole (13). This could be explained by the 3-5 times lower concentration of CDK4/6i that was used in our study compared to previous published work. Lower doses of drug were used in our experiments to avoid confounding biochemical changes in the cell as a result of toxicity or death processes. The exact mechanism of HLA upregulation and induction of antigen presentation by this class of inhibitors is still unknown. These results suggest that CDK4/6i has indirect effects on increased HLA surface expression, and a transcriptional program of antigen presentation does not seem to be induced, per se. Additionally, other studies showed induction of the immunopeptidome after treatment of MCF7 cells with Abemaciclib (14). As we also saw peptides derived from the immunoproteasome subunit PSMB10 presented on the cell surface after CDK4/6i treatment. This provided further evidence of the activation of the immunoproteasome that contributed to the re-shaping of the immunopeptidome.

Many of the drug-induced antigens we found on the CDK4/6i treated groups were associated with proteins from the mitotic pathways, specifically the G1/S phase transition, the phase in which cells arrest after CDK4/6i treatment (25). This suggested that inhibition of the cellular process led to preferential degradation of proteins and subsequent presentation on HLA peptides from proteins in the inhibited pathway. CDK4/6i gave rise to about 70 induced HLA ligands from the specific GO term “G1/S transition of mitotic cell cycle”, which represented about 10% of all treatment induced HLA ligands. Mass spectrometry analysis revealed the peptides “ALLESSLRQA” and “DRLVRAML” two HLA-A*02:01 and HLA-B*14:02 presented peptides from the Cyclin D1 protein, among other peptides. In a previous study the ALLESSLRQA sequence was predicted to be an HLA-A*02:01 binder, but had never been validated for its immunogenicity as it was considered a suboptimal HLA binder (26). By using RT-qPCR and Western blotting analysis we confirmed a compensatory upregulation of CDK4/6 binding partners, such as Cyclin D1, after CDK4/6i treatment. Additionally, Cyclin D1 exhibited higher levels of ubiquitination in CDK4/6i treated cells. Taken together, we propose a mechanism whereby disruption of the CDK4/CDK6/CCND1 complex leads to upregulation and subsequent degradation of cell cycle proteins (Figure 5). Similar changes in the immunopeptidome have also been reported, while this manuscript was in preparation, in melanoma cell lines after CDK4/6i treatment (27). Nonetheless, our study is the first to demonstrate pathway-dependent degradation and presentation.

Many of the drug-induced HLA ligands offer the potential to be ideal cancer targets for cancer immunotherapies. For example CCND1 is expressed in multiple cancers, e.g. mantle cell lymphoma (28) and native immunogenic peptides from this protein have been shown to induce CD4 and CD8+ cytotoxic T cell responses in HLA matched donors (29). Therefore, Cyclin D1 targeted immunotherapies may synergize with CDK4/6 inhibition in a variety of cancer types. Additionally, CCNE1 is a known negative prognostic marker in breast cancer (30) and an extensive study of HLA ligands in healthy tissues did not detect any peptides from CCNE1 presented on the cell surface of non-malignant cells (31). Given these characteristics CCND1 as well as CCNE1, together with the other induced CDK4/6i induced proteins, those targets should be considered promising targets for immunotherapy approaches.

CDK4/6i have considerable toxicity at clinically used antineoplastic therapeutic doses. Typically, CDK4/6i are administered in doses that result in plasma concentrations of 300-500nM, which are associated with hematologic toxicities including neutropenia and lymphopenia, as well as non-hematologic toxicities (i.e. diarrhea, nausea, dyspnea, arthralgia) (32). Our results suggested that using concentrations 3 to 5-fold lower than patient plasma concentrations can still achieve significant drug-induced peptide presentation changes. Using lower doses of CDK4/6i may mitigate toxicity while synergizing with immunotherapies (13).

Overall this study provides additional evidence and rationale to combine low-dose CDK4/6i with immunotherapies, such as checkpoint blockade, adoptive T cell transfer, TCRmimic antibodies or vaccination therapies. The robust induction of tumor specific proteins in conjunction with elevated overall HLA levels provide a basis for targeting these specific antigens in future immunotherapy-based combination regimens.

---

**Supplemental Figure 1.**
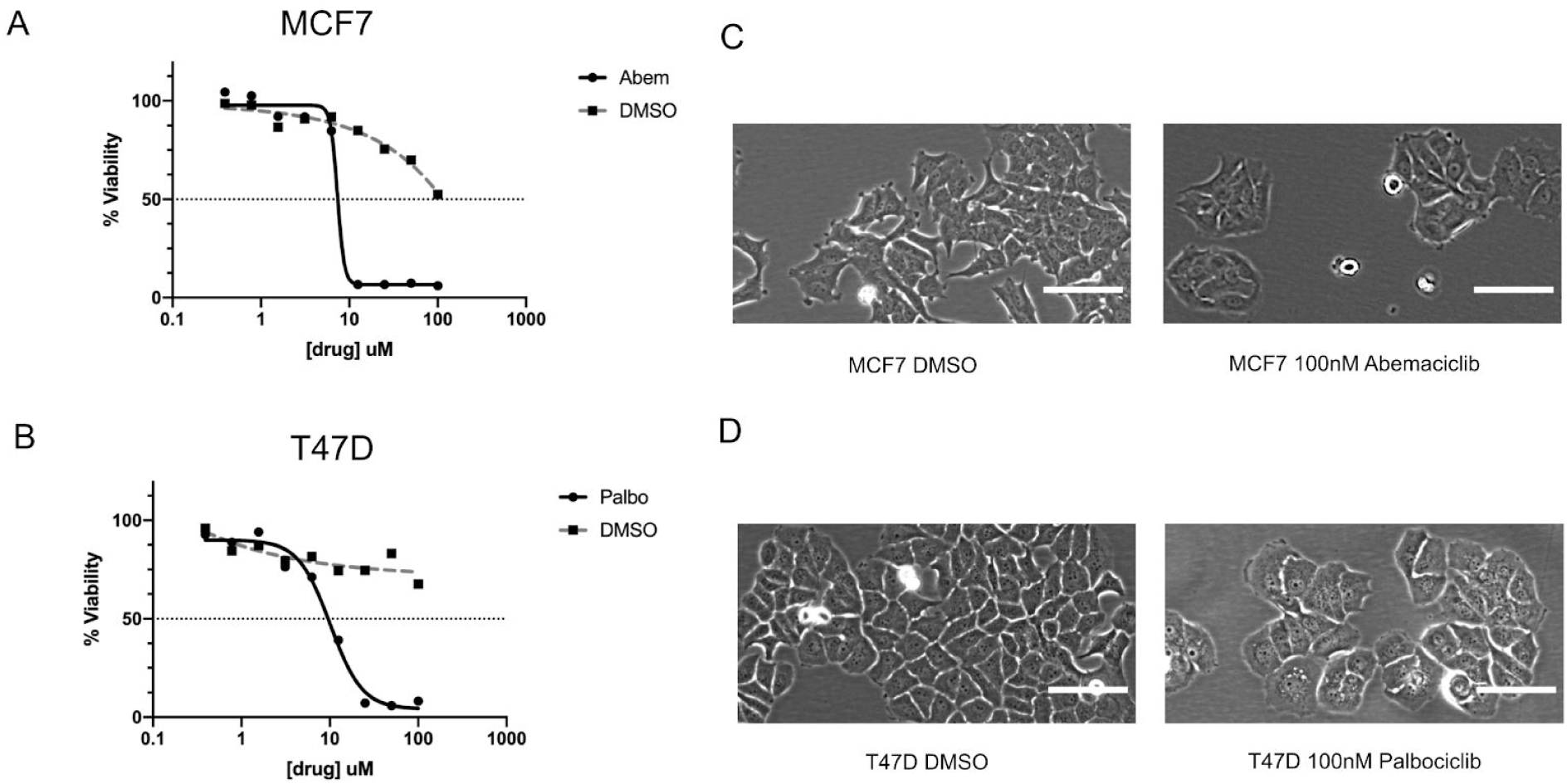
Low dose CDK4/6i upregulates HLA without toxicity. (A-B) Graph of cell viability of (A) MCF7 and (B) T47D cells treated with increasing doses of abemaciclib or palbociclib, respectively, as measured by Cell-Titer Glo at 72 hours. (C-D) MCF7 and T47D cells were treated with 100nM drug or DMSO as in (A) and bright field microscopy images were taken. Scale bars represent 100 pixels at the same magnification.

**Supplemental Figure 2.**
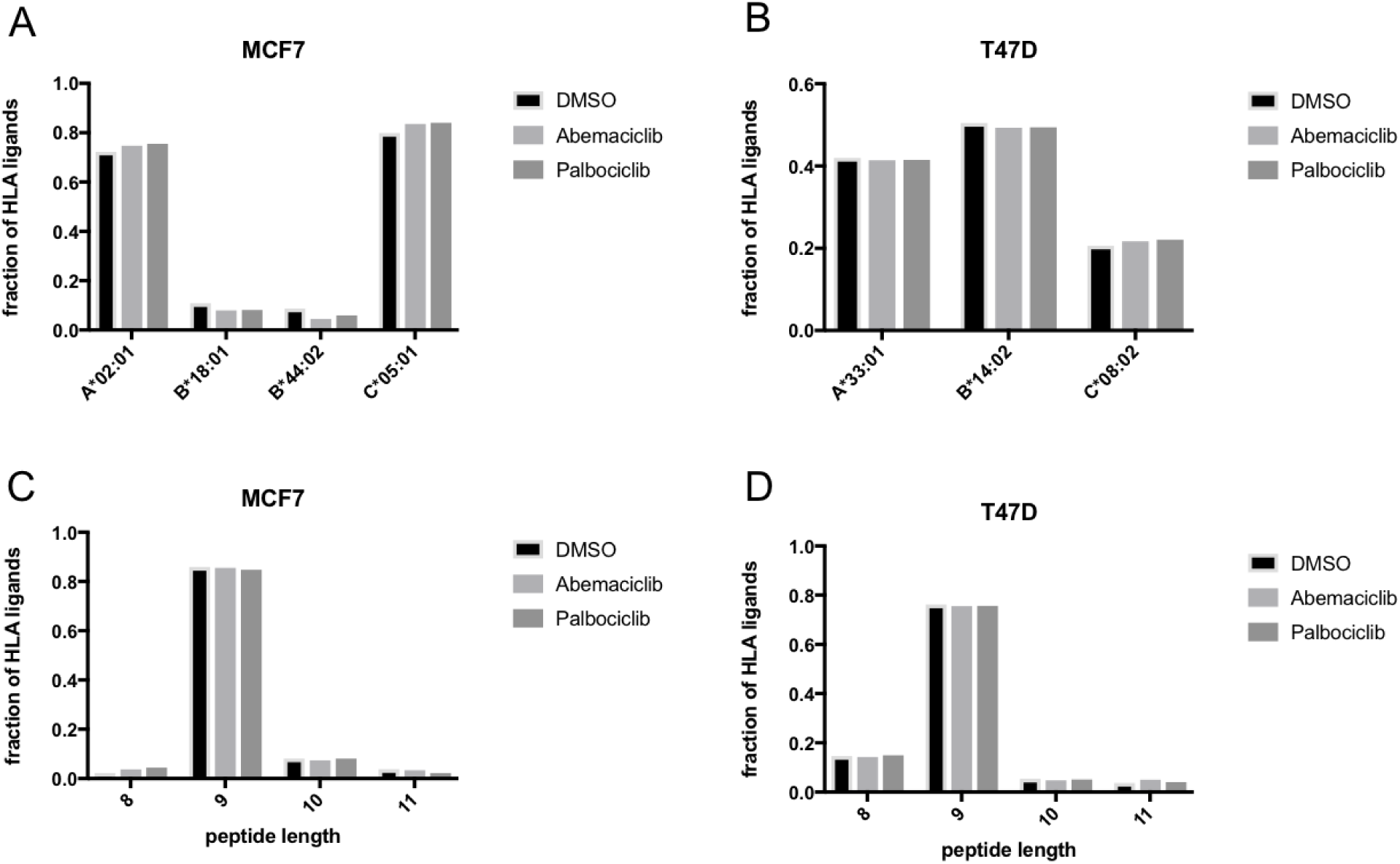
Characteristics of changes in the immunopeptidome after CDK4/6i treatment. Predicted binding preferences of HLA ligands after CDK4/6i treatment in (A) MCF7 and (B) T47D cells. Peptide length distribution of HLA ligands after CDK4/6i treatment in (C) MCF7 and (D) T47D cells.

## Notes

### Competing Interest Statement

D.A. Scheinberg has potential conflicts of interest, defined by please fill in Journal of interest by ownership in, income from, or research funds from: Pfizer, Sellas Life Sciences, Iovance, Eureka Therapeutics, and Bristol Myers Squibb.

## References

1. Fox P, Darley A, Furlong E, Miaskowski C, Patiraki E, Armes J, et al. The assessment and management of chemotherapy-related toxicities in patients with breast cancer, colorectal cancer, and Hodgkin’s and non-Hodgkin’s lymphomas: A scoping review [Internet]. European Journal of Oncology Nursing. 2017. page 63–82. Available from: http://dx.doi.org/10.1016/j.ejon.2016.12.008

2. Plummer C, Steingart RM, Jurczak W, Iakobishvili Z, Lyon AR, Plastaras JP, et al. Treatment specific toxicities: Hormones, antihormones, radiation therapy. Semin Oncol. 2019;46:414–20.

3. Sharpe AH, Pauken KE. The diverse functions of the PD1 inhibitory pathway [Internet]. Nature Reviews Immunology. 2018. Available from: http://dx.doi.org/10.1038/nri.2017.108

4. Ribas A, Wolchok JD. Cancer immunotherapy using checkpoint blockade. Science. 2018;359:1350–5.

5. Yin L, Dai S, Clayton G, Gao W, Wang Y, Kappler J, et al. Recognition of self and altered self by T cells in autoimmunity and allergy. Protein Cell. 2013;4:8–16.

6. Schumacher TN, Schreiber RD. Neoantigens in cancer immunotherapy. Science. 2015;348:69–74.

7. Dubrovsky L, Dao T, Gejman RS, Brea EJ, Chang AY, Oh CY, et al. T cell receptor mimic antibodies for cancer therapy. Oncoimmunology. 2016;5:e1049803.

8. Chandran SS, Klebanoff CA. T cell receptor-based cancer immunotherapy: Emerging efficacy and pathways of resistance. Immunol Rev. 2019;290:127–47.

9. Chang AY, Gejman RS, Brea EJ, Oh CY, Mathias MD, Pankov D, et al. Opportunities and challenges for TCR mimic antibodies in cancer therapy. Expert Opin Biol Ther. 2016;16:979–87.

10. Brea EJ, Oh CY, Manchado E, Budhu S, Gejman RS, Mo G, et al. Kinase Regulation of Human MHC Class I Molecule Expression on Cancer Cells. Cancer Immunol Res. 2016;4:936–47.

11. Oh CY, Klatt MG, Bourne C, Dao T, Dacek MM, Brea EJ, et al. ALK and RET Inhibitors Promote HLA Class I Antigen Presentation and Unmask New Antigens within the Tumor Immunopeptidome. Cancer Immunol Res. 2019;7:1984–97.

12. Planes-Laine G, Rochigneux P, Bertucci F, Chrétien A-S, Viens P, Sabatier R, et al. PD-1/PD-L1 Targeting in Breast Cancer: The First Clinical Evidences Are Emerging. A Literature Review. Cancers [Internet]. 2019;11. Available from: http://dx.doi.org/10.3390/cancers11071033

13. Goel S, DeCristo MJ, Watt AC, BrinJones H, Sceneay J, Li BB, et al. CDK4/6 inhibition triggers anti-tumour immunity. Nature. 2017;548:471–5.

14. Schaer DA, Beckmann RP, Dempsey JA, Huber L, Forest A, Amaladas N, et al. The CDK4/6 Inhibitor Abemaciclib Induces a T Cell Inflamed Tumor Microenvironment and Enhances the Efficacy of PD-L1 Checkpoint Blockade. Cell Rep. 2018;22:2978–94.

15. Mohammed AA, Rashied H, Elsayed FM. CDK4/6 inhibitors in advanced breast cancer, what is beyond? Oncol Rev. 2019;13:416.

16. Pernas S, Tolaney SM, Winer EP, Goel S. CDK4/6 inhibition in breast cancer: current practice and future directions. Ther Adv Med Oncol. 2018;10:1758835918786451.

17. Boegel S, Löwer M, Bukur T, Sahin U, Castle JC. A catalog of HLA type, HLA expression, and neo-epitope candidates in human cancer cell lines. Oncoimmunology. 2014;3:e954893.

18. Rozanov DV, Rozanov ND, Chiotti KE, Reddy A, Wilmarth PA, David LL, et al. MHC class I loaded ligands from breast cancer cell lines: A potential HLA-I-typed antigen collection. J Proteomics. 2018;176:13–23.

19. Loureiro J, Ploegh HL. Antigen presentation and the ubiquitin-proteasome system in host-pathogen interactions. Adv Immunol. 2006;92:225–305.

20. Vaddepally RK, Kharel P, Pandey R, Garje R, Chandra AB. Review of Indications of FDA-Approved Immune Checkpoint Inhibitors per NCCN Guidelines with the Level of Evidence. Cancers [Internet]. 2020;12. Available from: http://dx.doi.org/10.3390/cancers12030738

21. Gejman RS, Chang AY, Jones HF, DiKun K, Hakimi AA, Schietinger A, et al. Rejection of immunogenic tumor clones is limited by clonal fraction. Elife [Internet]. 2018;7. Available from: http://dx.doi.org/10.7554/eLife.41090

22. Wolf Y, Bartok O, Patkar S, Eli GB, Cohen S, Litchfield K, et al. UVB-Induced Tumor Heterogeneity Diminishes Immune Response in Melanoma. Cell. 2019;179:219–35.e21.

23. Heemskerk B, Liu K, Dudley ME, Johnson LA, Kaiser A, Downey S, et al. Adoptive cell therapy for patients with melanoma, using tumor-infiltrating lymphocytes genetically engineered to secrete interleukin-2. Hum Gene Ther. 2008;19:496–510.

24. Rafiq S, Purdon TJ, Daniyan AF, Koneru M, Dao T, Liu C, et al. Optimized T-cell receptor-mimic chimeric antigen receptor T cells directed toward the intracellular Wilms Tumor 1 antigen. Leukemia. 2017;31:1788–97.

25. Scott SC, Lee SS, Abraham J. Mechanisms of therapeutic CDK4/6 inhibition in breast cancer. Semin Oncol. 2017;44:385–94.

26. Chen J, Zurawski G, Zurawski S, Wang Z, Akagawa K, Oh S, et al. A novel vaccine for mantle cell lymphoma based on targeting cyclin D1 to dendritic cells via CD40. J Hematol Oncol. 2015;8:35.

27. Stopfer LE, Mesfin JM, Joughin BA, Lauffenburger DA, White FM. Multiplexed relative and absolute quantitative immunopeptidomics reveals MHC I repertoire alterations induced by CDK4/6 inhibition. Nat Commun. 2020;11:2760.

28. Wang M, Sun L, Qian J, Han X, Zhang L, Lin P, et al. Cyclin D1 as a universally expressed mantle cell lymphoma-associated tumor antigen for immunotherapy. Leukemia. 2009;23:1320–8.

29. von Bergwelt-Baildon MS, Shimabukuro-Vornhagen A, Wendtner CM, Kondo E. Identification of native, immunogenic peptides from Cyclin D1. Leukemia. 2010. page 209–11.

30. Zhao Z-M, Yost SE, Hutchinson KE, Li SM, Yuan Y-C, Noorbakhsh J, et al. CCNE1 amplification is associated with poor prognosis in patients with triple negative breast cancer. BMC Cancer. 2019;19:96.

31. Marcu A, Bichmann L, Kuchenbecker L, Backert L, Kowalewski DJ, Freudenmann LK, et al. The HLA Ligand Atlas. A resource of natural HLA ligands presented on benign tissues [Internet]. bioRxiv. 2019 [cited 2020 Apr 18]. page 778944. Available from: https://www.biorxiv.org/content/10.1101/778944v1

32. Choo JR-E, Lee S-C. CDK4-6 inhibitors in breast cancer: current status and future development. Expert Opin Drug Metab Toxicol. 2018;14:1123–38.

